# Integrating behaviour and ecology into global biodiversity conservation strategies

**DOI:** 10.1101/566406

**Authors:** Joseph A. Tobias, Alex L. Pigot

**Author notes:** Department of Life Sciences, Imperial College London, Silwood Park, Buckhurst Road, Ascot, Berkshire, SL5 7PY, UK.

## Abstract

Insights into animal behaviour play an increasingly central role in species-focused conservation practice. However, progress towards incorporating behaviour into regional or global conservation strategies has been far more limited, not least because standardised datasets of behavioural traits are generally lacking at wider taxonomic or spatial scales. Here we make use of the recent expansion of global datasets for birds to assess the prospects for including behavioural traits in systematic conservation priority-setting and monitoring programmes. Using IUCN Red List classification for >9500 bird species, we show that the incidence of threat can vary substantially across different behavioural syndromes, and that some types of behaviour—including particular foraging, mating and migration strategies—are significantly more threatened than others. When all factors are included in a combined model, behavioural traits have a weaker effect than well-established geographical and ecological factors, including range size, body mass and human population pressures. We also show that the association between behavior and extinction risk is partly driven by correlations with these underlying factors. Overall, these results suggest that a multi-species approach at the scale of communities, continents and ecosystems can be used to identify and monitor threatened behaviours, and to flag up cases of latent extinction risk, where threatened status may currently be underestimated. Our findings also highlight the importance of comprehensive standardized descriptive data for ecological and behavioural traits, and point the way forward to a deeper integration of behaviour into quantitative conservation assessments.

## 1. Introduction

Conservation biologists and behavioural ecologists have repeatedly called for closer links between their respective fields on the grounds that behavioural insights can contribute significantly to the success of conservation action (Clemmons & Buchholz 1997; Caro 1999; Caro & Sherman 2011; Greggor et al. 2016). However, this cross-disciplinary integration has progressed slowly, in part because the methods and central questions of behavioural ecology do not align closely with the needs of conservation practitioners (Greggor et al. 2016). For example, much of behavioural ecology focuses at the level of the individual, and identifies selective mechanisms acting on genes or organisms, whereas conservation typically operates at the level of populations (Caro 2007). This misalignment is perhaps most pronounced at macroecological scales where global analyses are playing a vital role in conservation science and policy (e.g. Newbold et al. 2015) but generally include only the most basic behavioural information.

One reason for the low profile of behaviour in comprehensive broad-scale analyses is because it is difficult and costly to measure standardised behavioural traits across species, space and time (Anthony & Blumstein 2000). The major contributions of behavioural research to conservation have dealt with factors such as individual movements, sensory ecology or animal personality, and the extent to which they mediate various kinds of human pressures, including disturbance, habitat loss and hunting (Greggor et al. 2016). The key behavioural metrics under this framework are context-dependent, highly plastic both within and between individuals, and typically estimated through detailed observation and experimentation. They are often inappropriate for quantitative assessments at the wider level of communities or ecosystems because they are (1) only available for a small fraction of species, and (2) not readily incorporated into species-level analyses. For instance, the case-dependent intricacies of how behaviour influences Effective population size (*N*_e_) are useful to conservation (Anthony & Blumstein 2000) but we are decades away from having these data available for comprehensive global studies.

Global or regional conservation assessments are largely restricted to comprehensive species-level datasets accessible at the relevant scale (see figure 1). Most macroecological analyses have therefore tested whether species conservation status is predicted by human impacts, biogeographical factors such as latitude or range size, and environmental factors such as climate or habitat (Bennett & Owens 1997, Owens & Bennett 2000, Cardillo et al. 2004, Cardillo et al. 2005, Lee & Jetz 2011, Keinath et al. 2017), or reversed the process to predict the conservation status of poorly known species (Jetz & Freckleton 2015, Santini et al. 2019). Using freely available GIS layers, these socio-economic, biogeographical and environmental variables can be extracted for specimen localities or geographical range polygons, which in some vertebrate groups are reasonably accurate. The other main components of macro-scale assessments have been demographic factors, including population size and density, and rates of population decline, all of which are theoretically related to extinction risk (Keinath et al. 2017; Santini et al. 2019). In general, only crude population estimates are included in global-scale analyses because very few attempts have been made to quantify population sizes and trends across entire global ranges (Tobias & Seddon 2002, Tobias & Brightsmith 2007). Previous studies have shown that both extrinsic biogeographic and demographic factors are correlated with extinction risk, leading to their widespread inclusion in regional and international conservation status assessments.

**Figure 1.**
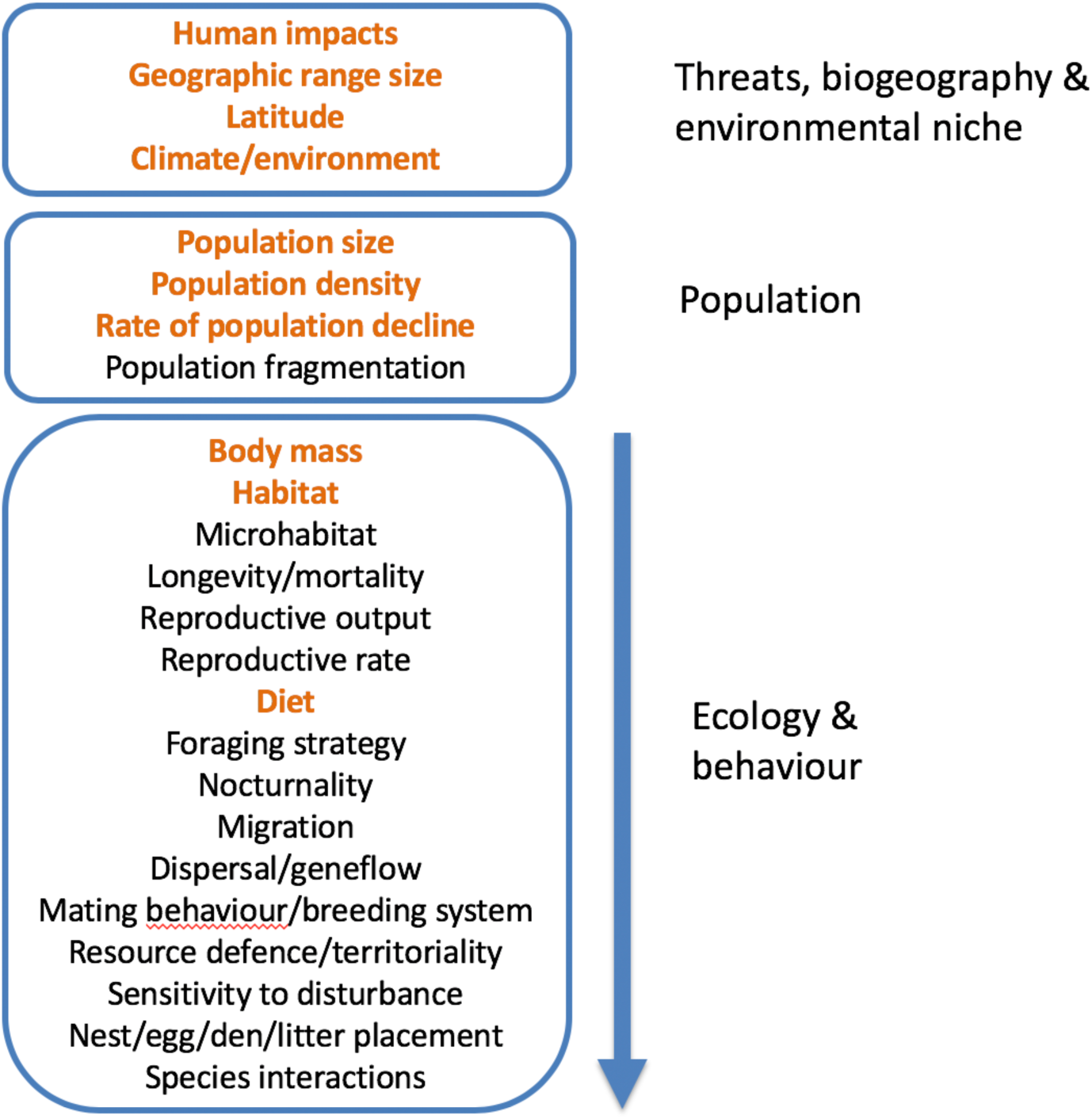
Extrinsic and intrinsic factors associated with extinction risk or conservation status at global scales. Extrinsic factors include anthropogenic threats to species and the biogeographic and environmental context; intrinsic factors include population and ecological niche dimensions. This diagram summarises the types of traits that are either available or desirable when constructing models of conservation risk at macroecological (continental or global) scales; numerous additional factors may impinge on conservation assessments in particular clades or species. Red text indicates datasets currently available for all species in well studied groups like birds. Availability of data is currently biased towards environmental, biogeographical and population attributes, whereas data tend to be unavailable, uncertain or sparse for most ecological variables, and absent for behavioural variables.

Perhaps the most influential global assessment is the IUCN Red List (IUCN 2001), an indicator of biodiversity status and change linked to international convention targets (Butchart et al. 2005). The conservation status categories systematically generated by the Red Listing process are enshrined in legislation and widely used in macroecological research (Rodrigues et al. 2006). Previous assessments of predictors of Red List status have generally focused on standard biogeographic or climatic variables, without delving far into behavioural or ecological factors. Indeed, the only ecological and behavioural traits incorporated into most global models of conservation risk are body mass, diet and habitat preferences (Lee & Jetz 2011; Newbold et al. 2015, Keinath et al. 2017). To convert these variables into species-level traits, body mass is typically averaged from small numbers of published estimates, while diet and habitat are classified into broad categories on the basis of published descriptions in secondary literature (Wilman et al. 2014). By contrast, the availability of many other behavioural or ecological variables is highly patchy at global scales, and limited by the difficulty of converting into species-level traits (figure 1).

The most relevant behavioural traits to conservation assessment include those that mediate sensitivity to habitat loss, fragmentation, and climate change (Greggor et al. 2016). Factors relating to dispersal behaviour are particularly pertinent because they impinge on the ability of species to cross unsuitable habitat and thus maintain interconnected metapopulations after habitat fragmentation (Lees & Peres 2009). Dispersal-related traits may also regulate the ability of species to track shifting geographical ranges in response to climate change (Early & Sax 2011, Howard et al. 2018), and predict susceptibility to threats like wind farms (Thaxter et al. 2017). In addition, behavioural dimensions of species interactions may be important determinants of responses to a variety of threats. For example, studies focused at the level of species pairs or communities find evidence that interspecific competition leads to population declines or local extinction following habitat loss and fragmentation (Bregman et al. 2015, Grether et al. 2017) while reproductive interference may threaten populations of closely related species interacting or hybridising when climate-driven range shifts lead to secondary contact (Hochkirch et al. 2007, Greggor et al. 2016). However, while standardised estimates of dispersal ability and interspecific competition are available for restricted samples of species, they are not readily available at macroecological scales, except in the form of extremely coarse categories (e.g. whether an organism can fly or not; Keinath et al. 2017).

Other variables potentially relevant to conservation status can be placed on a continuum from primarily ecological to primarily behavioural (figure 1). At the ecological end are aspects such as microhabitat preferences, while other factors such as foraging mode, migration, sexual selection, territoriality, reproductive strategy and nesting behaviour have an increasingly behavioural dimension. Previous research suggests that species sensitivity to land-use or climate change can be related to microhabitat (e.g. in the form of vertical stratum of vegetation), foraging behaviour (e.g. gregarious foraging), and reproductive strategy (e.g. breeding system) (Kokko & Brooks 2003, Bueno et al. 2018). Similarly, territorial strategy is linked to species sensitivity to habitat fragmentation (Ulrich et al. 2017), suggesting that elevated interspecific competition via behavioural mechanisms can increase threats associated with land-use and climate change (Jankowski et al. 2011, Grether et al. 2017). Until recently, such inferences were based on relatively restricted species sampling, but this constraint is changing as the compilation and dissemination of global trait datasets gathers pace.

To assess whether recent progress in data availability can pave the way for behavioral perspectives to be explicitly included in global conservation strategies, we compiled information on a variety of ecological and behavioural traits for all bird species, including estimates of sexual selection (Dale et al. 2015; Cooney et al. 2017), breeding system (Jetz & Rubenstein 2011), foraging strategy (Pigot et al. 2016, Felice et al. 2019), territorial behaviour (Tobias et al. 2016), and nest placement (Stoddard et al. 2017). We then ran multivariate models to evaluate the extent to which behaviour predicts IUCN Red List status at macroecological scales and in relation to a range of standard biogeographical and environmental variables. Our goal is to assess the current landscape of behavioural data availability and the prospects for more nuanced conservation assessments and priority-setting.

## 2. Methods

### (a) Data

We assembled data on species threat status from the 2016 Red List (IUCN 2016) along with a range of potential drivers of variation in status, including biogeographic, ecological and behavioural traits, as well as the exposure of each species to human impacts. Geographic range size is consistently identified as the strongest predictor of threat status (Lee & Jetz 2011; Jetz & Freckleton 2015). We estimated range size for each species based on expert opinion extent of occurrence maps of species breeding distributions (BirdLife International, 2012). Human population pressure is also known to influence extinction risk (Cardillo et al. 2004; Scharlemann et al. 2005; Davies et al. 2006). To quantify the exposure of species to human impacts, we first extracted polygon range maps onto an equal area grid (resolution of 110 km ≈ 1° at the equator) and used this grid to sample human population density, human appropriation of net primary productivity and night-time light intensity, an indicator of urbanisation and development. We calculated the mean value of each metric, averaged across all grid cells overlapping with each species range.

We collated data on a selection of ecological traits, including mean species body mass (g), habitat type, diet and island dwelling, all of which have been linked to extinction risk (Bennett & Owens 1997; Owens & Bennett 2000; Cardillo et al. 2005; Lee & Jetz 2011; Jetz & Freckleton 2015). We assigned species to one of ten dietary categories: aquatic animals, aquatic plants, terrestrial invertebrates, terrestrial vertebrates, terrestrial carrion, nectar, seeds, fruit, other terrestrial plant matter (e.g. leaves) and omnivore, based on the dominant resource present in their diet (see Supplementary material). Data on proportional resource use were first obtained from Wilman et al. (2014), and then modified and updated based on comprehensive literature searches. Our dietary classification differs from Wilman et al. (2014) in that we subdivided each animal or plant-based resource type into separate aquatic and terrestrial categories (see Felice et al. 2019). This helps us to avoid highly heterogenous categories such as invertivores, which spans a wide variety of species from insectivorous warblers to squid-eating albatrosses and crustacean-eating flamingos (Wilman et al. 2014). Our approach separates warblers (diet: “terrestrial invertebrates”) into a different category from albatrosses and flamingos (diet: “aquatic animals”). Using literature to score habitat use, we assigned species to broad habitat categories (coastal, terrestrial, freshwater, sea) according to the predominant habitat utilised across their geographic distribution. We included habitat type as a predictor in our main models but also used this variable along with a measure of forest dependency (obtained from BirdLife International: http://datazone.birdlife.org/home) to subset our data and perform additional analysis focusing on terrestrial species (n = 8433) or those with medium to high forest dependency (n = 5646). Using the geographical range polygons described above, we classified species as island dwelling if more than 25% of their geographic range occurred on small islands (landmass <2000 km2). Further details of data compilation methods are given in supplementary materials.

To assess the association between IUCN threat status and key behavioural traits, we assembled data on foraging strategy, nest placement, breeding system, mating behaviour, the mean clutch size of broods, territoriality and migratory behaviour. Following the method described by Felice et al. (2019), we used literature searches to assign species to one of seven foraging strategies. We classified each species according to the predominant behavioural strategy used to acquire resources, and assigned species utilising multiple foraging strategies as generalists (see Supplementary material). Nest placement was scored into a simple three-way system: ground, elevated or cavity (see Stoddard et al. 2017 for details). We used a binary score of breeding system based on a published classification of cooperative and noncooperative breeders (Jetz & Rubenstein 2011). Mating behaviour was scored as strict monogamy, monogamy with infrequent (<5% males) polygyny, monogamy with frequent (5-20% males) polygyny, and polygamy (>20% males and females). These categories are based on the index of sexual selection developed by Dale et al. (2015). Clutch size data was based on Jetz et al. (2008). Using data from Tobias et al. (2016), we assigned all species to three categories according to the degree of territoriality: ‘strong’ (territories maintained throughout year), ‘weak’ (weak or seasonal territoriality, including species with broadly overlapping home ranges or habitually joining mixed species flocks), and ‘none’ (never territorial or at most defending very small areas around nest sites). Finally, we assigned the migratory behaviour of species as either sedentary, partially migratory (minority of population migrates long distance or most individuals migrate short distances) and migratory (majority of population undertakes long-distance migration) (Tobias et al. 2016).

Most variables were available for the vast majority (i.e.>99%) of species but the identity of species with missing values differed across variables. For categorical predictors, we imputed missing values using the modal class for each genus, if the genus contained at least 2 species and the modal class was present across at least 75% of species. If these conditions were not met, we used the same criteria to either impute missing values at the family level. After removing all species with any missing values, our final dataset included n = 9576 species.

### (b) Statistical analysis

To model the effects of each predictor variable on extinction risk, we treated threat as a binary variable (0, 1) according to the IUCN Red List categories. All species listed as Vulnerable, Endangered, Critically Endangered, Extinct (including Extinct in the Wild) were classified as Threatened; the remainder (Near Threatened, Least Concern and Data Deficient) were classified as non-Threatened. We modelled threat using a generalised linear mixed effects model, with a binomial error structure and including taxonomic family as a random effect to control for the phylogenetic non-independence of species when identifying predictors of threat. Predictor variables exhibiting right skew were log transformed prior to analysis.

In contrast to previous assessments of the predictors of extinction risk in birds (e.g. Lee & Jetz 2011), we are particularly interested in how behaviour and its covariation with other putative drivers of extinction risk alter the incidence of threat. First, to assess the overall association between each predictor and threat, we fitted a series of single predictor (i.e. univariate) models. Second, we generated a series of multivariate models and calculated relative model fit according the Akaike Information Criterion (AIC). We assessed the relative importance of each behavioural trait relative to other predictors by excluding each variable in turn from the full model and calculating the difference in AIC (delta AIC). We then assessed the overall contribution of behavioural traits in predicting threat by calculating the AIC and r^2^ of a full model including all predictor variables, and comparing with a model excluding behavioural traits. Finally, we also calculated the AIC and r^2^ of a model including only behavioural traits.

When comparing univariate and multivariate models, we were particularly interested in testing how covariation between behavioural traits and ecological, geographical or socio-economic variables may modify the association between threat and behaviour. We identify three possible scenarios. First, when behaviour is weakly related to threat, we may nevertheless find strong variation in the incidence of threat across behavioural categories because of differences in other factors that drive variation in threat (i.e. ecology, geography or human impacts), an example of an ‘enhanced’ effect. Second, the opposite pattern may emerge, if behaviour has a significant effect on threat, but this effect is ‘masked’ by countervailing effects of ecological, geographical or human impacts. Finally, the apparent effect of a given behaviour on threat could even be ‘reversed’, when taking into account covariation with other factors.

To examine how the definition of threat may influence the predictors of extinction risk, we repeated our analysis considering only threatened species (*n* = 1216), predicting lower (0 [Vulnerable]) or higher (1 [Endangered, Critically Endangered, Extinct]) levels of threat. To assess how the predictors of threat may change across broad habitat types, we repeated our analysis on different subsets of our data including all species (n = 9576), terrestrial species (n = 8433) and forest dependent species (n = 5646).

## Results

Our results identified a number of core predictors of threat status that align closely with previous assessments indicting that threat arises as a combination of geography, ecology and human impacts (Fig. 2). Specifically, the strongest predictor of threat status is geographical range size, with additional strong effects of body mass, island dwelling and the mean human population density across the species geographic range, a metric of exposure to human impact. In both univariate and multivariate models, the incidence of threat decreases with geographic range size and increases with body size (Table S1). When tested in isolation, the incidence of threat is higher on islands than on the mainland and in areas of low human population density (Table S1). However, in the full multivariate model, these effects are reversed, with a higher incidence of threat in areas of greater human population density, but a lower incidence of threat on islands (Table S1, see also Manne et al 1999). In addition to these core predictors, we also identified an effect of behaviour on extinction risk. A multivariate model including behavioural traits alone explains 7% of the variance in threat status. A full multivariate model including all predictors is significantly better supported than one excluding behavioural traits (delta AIC = 36) although the improvement in explanatory power is small (R^2^ excluding versus including behaviour = 0.49 versus 0.51 respectively). Behavioural traits receiving strong support for inclusion in the full multivariate model (delta AIC >2) were mating behaviour (monogamous or polygamous mating) and migration (Fig. 2). Behavioural traits receiving weak or no support for inclusion in the model (delta AIC <2) were foraging mode, breeding system, territoriality, nest placement and clutch size. The effects of behaviour were similar regardless of whether we conducted our analysis across all birds, only terrestrial species (Fig. S1a) or those restricted to forests (Fig. S1b). This is perhaps not surprising, given that forest dependent species comprise more than half of all birds. We note, however, that the role of behaviour in predicting threat does depend on the way in which threat is defined. Specifically, while behaviour is a significant predictor of whether a species is threatened or not (delta AIC = 36), it does not predict the level of threat (i.e. whether a species is Vulnerable versus Endangered, Critically endangered or Extinct) (delta AIC = −15, Fig. S2).

**Figure 2.**
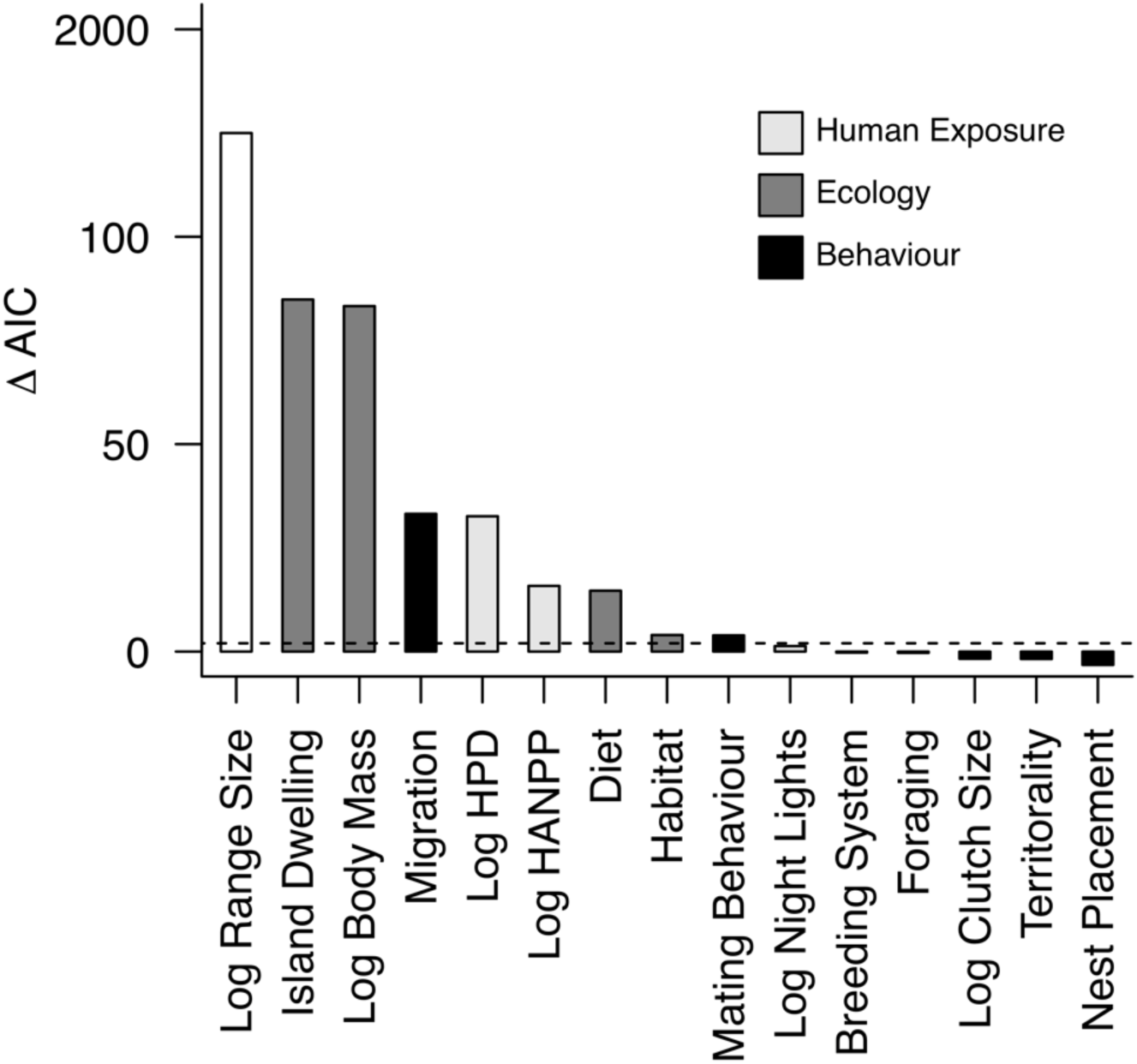
The relative contribution of anthropogenic, ecological and behavioural variables to explaining threat status. Variable contributions are quantified as the difference in AIC between the full model and a model excluding each variable. Variables are colored according to variable type. The dashed line indicates a difference of 2 AIC units indicating strong support for variable inclusion.

Some behavioural traits were unrelated to threat, regardless of whether they were considered in isolation or in the full multivariate model. In particular, we found no effect of nest placement or breeding system in our models (Fig. 3, Table S1). In other cases, threat exhibited significant associations with behaviour, but with effects that varied depending on whether we accounted for other putative drivers of extinction risk (Fig. 3a, Table S1). In the case of foraging behaviour, we find that the incidence of threat varies substantially across foraging categories. For instance, >30% species that feed either by diving or by aerial attacks in aquatic habitats are threatened compared to <10% species that are foraging generalists or bark probing specialists in terrestrial habitats (Fig. 3a). However, our full multivariate model shows that most of this variation in the incidence of threat is driven by covariation between foraging behaviours, ecological traits and exposure to human impacts (Fig. 4). In particular, species feeding at sea and with large body size are more threatened than land-based and small bodied species (Fig. S3). Having accounted for these confounding variables, only species feeding by aerial attacks in aquatic habitats have significantly higher levels of threat (i.e. an example of an ‘enhanced’ effect) (Fig. 4, Table S1). A similar effect was also found for clutch size (Table S1). While the incidence of threat declines with increasing clutch size, this association is not significantly supported when accounting for confounding variables in the full multivariate model (Table S1).

**Figure 3.**
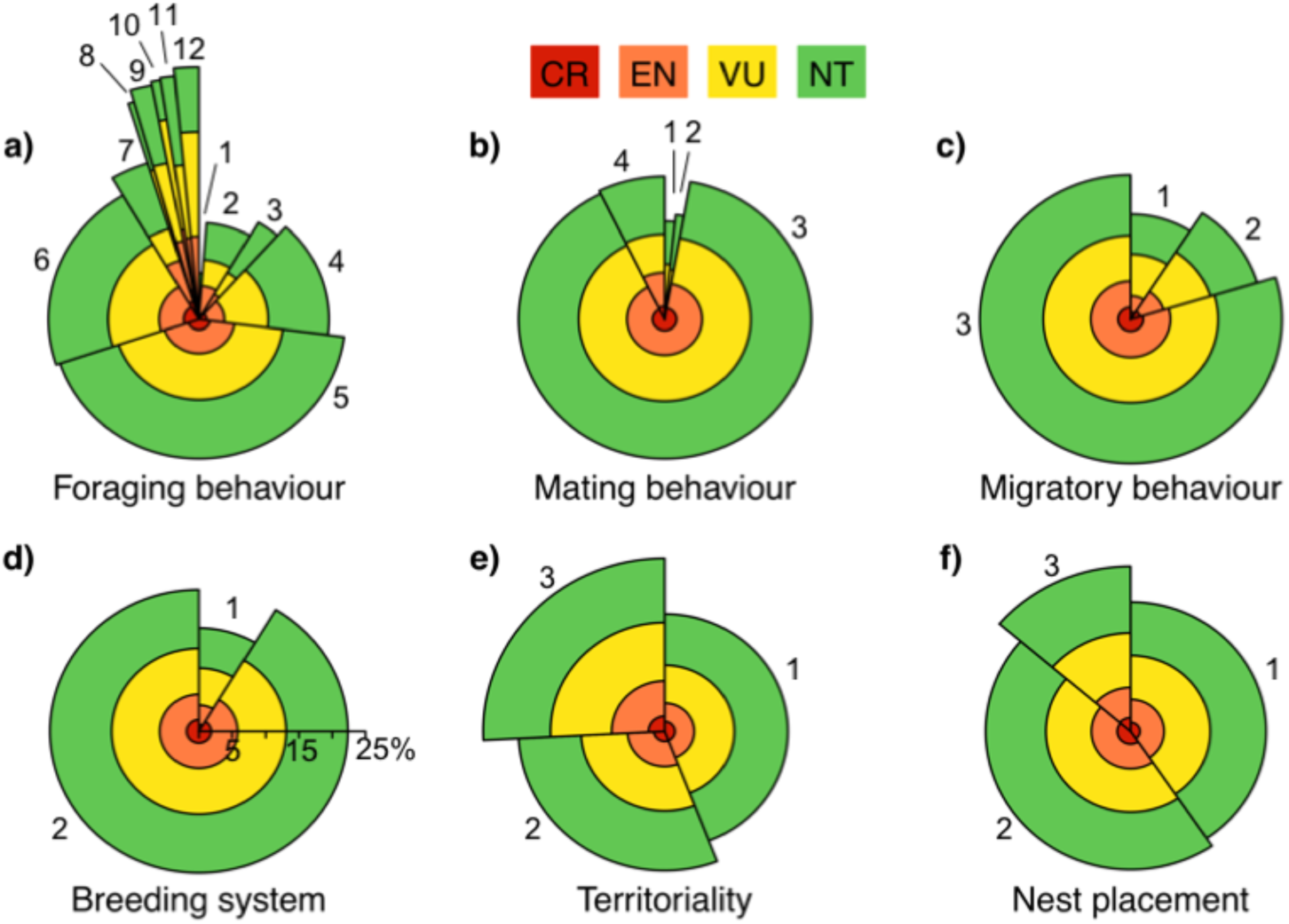
The % of threatened species in different behavioural syndromes: a) Foraging behaviour (1 Foraging generalist, 2 Aerial screen, 3 Bark glean, 4 Aerial sally, 5 Arboreal glean, 6 Ground forage, 7 Aquatic ground, 8 Aquatic plunge, 9 Aquatic surface, 10 Aquatic aerial, 11 Aquatic generalist, 12 Aquatic dive), b) Mating behaviour (1 Monogamy with infrequent polygyny, 2 Monogamy with frequent polygyny, 3 Monogamy, 4 Polygyny), c) Migratory behaviour (1 Migrant, 2 Partial migrant, 3 Sedentary), d) Breeding system (1 Cooperative, 2 Non-cooperative), e) Territoriality (1 Weak, 2 Strong, 3 None), f) Nest placement (1 Cavity, 2 Exposed elevated, 3 Exposed ground). The width of each segment indicates the proportion of all species (N = 9576) in each behavioural syndrome. Segment heights indicate the % of species that are threatened in each syndrome. Colours indicate threat level (Critically endangered [CR], Endangered [EN], Vulnerable [VU] and Near Threatened [NT]).

**Figure 4.**
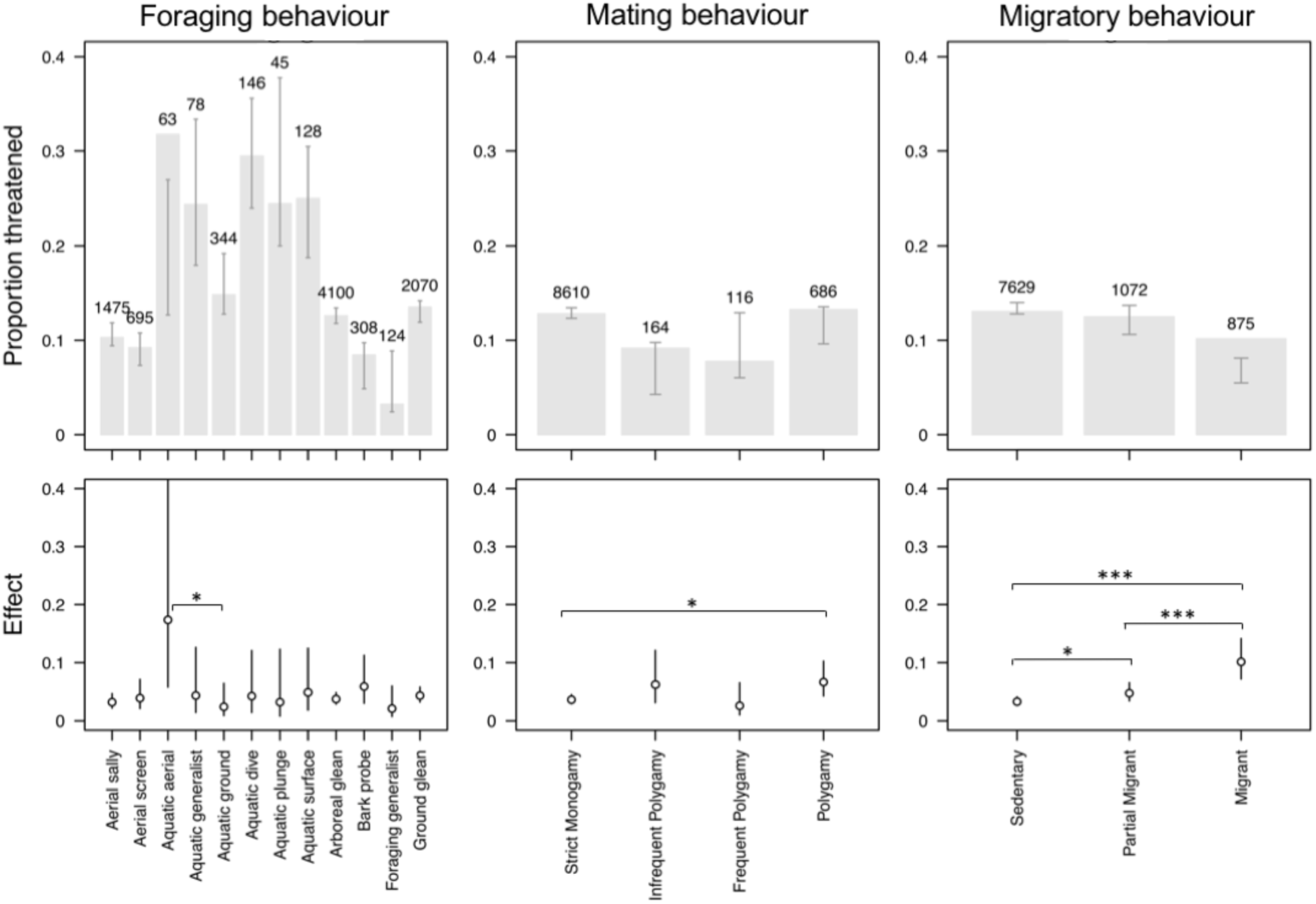
Variation in the prevalence of threat across behavioural syndromes. Top row: bars indicate the observed proportion of threatened species for different foraging modes, mating behavioural and migratory strategies. Brackets indicate the expected proportion (95% CI) of species threatened based on all other predictor variables. Numbers indicate the number of species in each category. Bottom row: the estimated effect size of each behavioral category (mean and 95% CI). Significant contrasts are indicated at the *p* = 0.05 (*), *p* = 0.01 (**) and *p* = 0.001 (***) level. Effect sizes and significance was assessed with a generalized linear mixed effects model including all predictor variables and family as a random effect.

Mating behaviour provides a possible example of a ‘masking’ effect. When tested in isolation, we found that polygamous species are no more likely to be threatened than monogamous species (Fig. 3b, Table S1). However, after accounting for the confounding effects of other predictors in the full multivariate model, we found that the probability of being threatened is significantly higher among polygamous than monogamous species (Fig. 4, Table S1). This effect of mating behaviour is masked when considered in isolation because polygamous species on average have a smaller body size than monogamous species, and this smaller body size nullifies the effect of mating behaviour on threat (Fig. S3). This suggests that polygamy may enhance the risk of extinction but that its effects may have been masked due to covariation with other factors that decrease extinction risk.

In a univariate model, we found that proportionately fewer migrants are threatened compared to partial migrants or sedentary species (Fig. 3c, Table S1). This may suggest that migration, or perhaps associated greater vagility, buffers species from extinction. However, in our full model we found the opposite effect of migration on the likelihood of being threatened, whereby migrants are more likely to be threatened than sedentary species (Fig. 4). These contrasting findings arise because migratory tendency is strongly correlated with range size, with migrants have larger breeding ranges on average than sedentary species (Fig. S3), and thus a lower incidence of threat. However, having statistically accounted for the negative effects of range size on threat, migrants are more likely to be threatened than sedentary species (Fig. 4, Table S1). This suggests that undertaking long distance migration makes species more at risk of extinction but that this is unable to overcome the effects of other covarying factors that instead lead to a higher prevalence of threat among sedentary species (i.e. an example of a ‘reversed’ effect).

## Discussion

We have shown that global-scale ecological and behavioural datasets predict variation in IUCN Red List status of birds, but that these relationships are largely explained by underlying correlations with well-established macroecological variables. Some behavioural traits were only significant predictors when behaviour was analysed independently, becoming non-significant when correlations with factors such as body size, geographical range size and human impacts were included. Conversely, other behavioural traits were not significant predictors in behaviour-only models, and their effect was only evident when socio-economic and biogeographic variables were included. These findings are consistent with previous reports that most ecological and behavioural traits have relatively weak associations with conservation status when incorporated into regional or global models as a species-level trait (Lee and Jetz 2011, Newbold et al. 2015, Keinath et al. 2017). However, although we find little evidence that the recent expansion of behavioural datasets can contribute substantially to refining conservation strategies at these wider scales, our results also show that behavioural traits act as modifiers that can improve explanatory power in conservation assessments and other predictive exercises.

The traits with strongest influence on conservation status were foraging strategy, mating behaviour and migration. Even in these cases, we found that significant relationships between behaviour and conservation status were only detected for certain strategies. For example, bird species foraging by diving from air to water were significantly more threatened than otherwise predicted. Moreover, a number of species-level behaviours, including variation in breeding system, territoriality, and nest placement, had little predictive power in explaining variation in IUCN Red List status regardless of how they were entered into models. This does not necessarily indicate that such factors are unimportant to conservation, as it is well known that they play a role in some contexts (e.g. nest design and placement has important implications for predation risk in modified landscapes; Wilcove 1985). However, our models show that these effects are minor and often overwhelmed by other non-behavioural factors at global scales.

These results do not support the integration of behaviour into global conservation assessment frameworks, including the IUCN Red List criteria. However, the accuracy of Red List assessments might be improved by using life history and behaviour to scale terms in the criteria which are difficult to assess or define, such as “number of mature individuals” and “future rate of decline” (IUCN 2001). These factors are typically judged with a large dose of guesswork (see Tobias & Seddon 2002, Tobias & Brightsmith 2007). Guidelines on how to scale judgements in relation to ecological and behavioural factors such as mating systems, sex ratios, reproductive rate and predation pressure could be useful in fine-tuning predictions of “number of mature individuals” and “future rate of decline”. Moreover, for Red List assessors considering what constitutes “severe fragmentation”, future versions of the criteria may be improved with guidelines on how best to account for dispersal ability, gap-crossing ability and ecological specialism.

### (a) Challenges

Our findings highlight one of the key challenges of applying behavioural data over larger spatial and taxonomic scales, namely that behavioural traits can have a major influence in particular species or contexts, yet only reduced effect in global analyses. This occurs for two main reasons. First, behavioural traits are inherently flexible within and between individuals and therefore poorly represented by averaging across entire species or populations. Second, behaviour is often not consistently or independently associated with extinction risk in the same way as, for example, low population size, small geographic range and slow reproductive output (Cardillo 2005, Lee and Jetz 2011).

This point can be illustrated by year-round territoriality, a system of resource defence most widespread in tropical birds (Tobias et al. 2016). Intense year-round territorial behaviour can increase the risk of extinction in some contexts, such as mountaintop species driven to extinction through costly agonistic interactions with lower elevation replacements moving upslope in response to climatic warming (Jankowski et al. 2011, Freeman et al. 2018). The costs of territoriality are asymmetric, producing both lower-elevation winners and upper-elevation losers. Moreover, the pattern of non-overlapping elevational ranges for highly territorial species holds largely true for some species pairs and localities (Freeman et al. 2019), but not others (Boyce & Martin 2019), particularly in lowland systems where species do not tend to occupy rare climatic niches or to share parapatric range boundaries with close ecological competitors. Given that the relationship between territoriality and extinction risk is bidirectional and context-dependent, it makes sense that we find no overall link between territoriality and IUCN Red List status.

An important viewpoint to bear in mind is that the models presented here treat behaviour as an independent species-level trait whereas the influence of behaviour is often dependent on inter-relationships among species. Staying with the example of territoriality, the key factor is not so much whether a particular species aggressively defends territories year-round, but whether it directly competes with a closely related taxon that does the same. Thus, future versions of global models or associated conservation assessments should consider scoring behavioural interactions rather than behaviour per se. Advancing towards this goal is particularly urgent given that species interactions are sensitive to environmental effects. Both climate and land-use change can potentially influence the behavior of multiple interacting species, as well as their phenology, physiology and relative abundance, and we ideally need to quantify a range of behavioural interactions and responses to understand how environmental changes affect interaction-based ecosystems (Tylianakis et al., 2008). Again, the key challenge is that the role of behavior in heterotrophic systems can be complex and highly flexible (Ness & Bressmer 2005), creating difficulties for multi-species models. Nonetheless, we may improve predictions by incorporating behaviour in more sophisticated ways using interaction-based models, starting at local scales and expanding to larger scale ecological networks when data become available.

A related point is that, although we have largely focused on how particular behaviours may influence extinction risk, such factors may yet prove to be less important than behavioural flexibility itself (Sol et al. 2016). Individual organisms with the ability to modify their behaviour through adaptability (i.e. plasticity) may be better able to survive when confronted with novel environmental conditions and selection pressures imposed by anthropogenic change. Defining and developing general indices of behavioural flexibility and innovation remains a challenge (Audet & Lefebvre 2017), but may nevertheless be broadly predictable by morphometric traits that are increasingly available at large scales (Sol et al. 2005). For instance, differences in relative brain size across species is positively associated with rates of behavioural innovation in birds, an effect that may explain the apparently greater success of large brained species in colonising and persisting in more unpredictable environments (Sayol et al. 2006, Sol et al. 2008), including cities, the most highly altered of human environments (i.e. the ‘cognitive buffer’ hypothesis) (Sol et al. 2013).

### (b) Opportunities

Although they extend the number of behavioural traits compiled across a major global radiation, our analyses are limited by the patchy availability of trait datasets and thus remain highly incomplete (figure 1). A major omission is dispersal behaviour, which we only include as a simple score of migration. Dispersal has long been considered relevant to the conservation of fragmented populations and the optimum design of reserve networks (Caro 1999). However, despite the likely importance of dispersal to understanding biodiversity responses to habitat loss and fragmentation, most broad-scale models (e.g. Newbold et al. 2013, Bregman et al. 2014) lack estimates of dispersal behaviour simply because they are generally not available as a standardised organismal trait at macroecological scales. This problem may be addressed by the fast-moving field of movement ecology, with GPS trackers and loggers deployed over increasing numbers of species (Kays et al. 2015), and data compilation accelerated by new satellite tracking systems, such as ICARUS (https://icarusinitiative.org). Given that it could take decades for these technological innovations to generate comprehensive dispersal estimates across major taxonomic groups, one potential stopgap solution is to use morphometric indices of dispersal or flight ability. Dispersal indices, such as hand-wing index in birds, can be estimated by measuring museum specimens to provide a fuller picture of spatial ecology and movement behaviour across multiple species in macroecological analyses (e.g. Pigot & Tobias 2015) and comparative studies of anthropogenic threats (e.g. Thaxter et al. 2017). Such indices, along with further missing data on factors such as reproductive rate and sensitivity to disturbance (figure 1) should be compiled and applied to conservation assessments at global scales.

Another area where behavioural indices may prove useful is ecological forecasting. At present, dispersal is usually ignored in global range shift models, or only included on the basis of extremely crude metrics, such as geographical range size (e.g. Hof *et al.* 2018). Similarly, species interactions are difficult to quantify and, while most range shift forecasting models acknowledge the limitation, they are generally not included in analyses. Future models should explore the possibility of estimating the strength of species interactions using either pairwise morphometric trait divergence or scores of territorial behaviour, both of which have been shown to limit geographical range overlap in pairs of avian sister species (Pigot & Tobias 2013, Freeman et al. 2019). Theoretically, suites of behavioural traits and associated morphometric indices can be incorporated into species distribution modelling in much the same way proposed for detailed physiological traits (Chown 2012).

The associations we detect between behaviour and conservation status (figure 3) suggests that future research could use similar techniques to identify “threatened behaviours” or suites of behaviours. Using global analyses to look beyond species conservation and instead to identify behaviours that are rare or declining might be a useful step towards targeting conservation action towards maintaining behavioural trait diversity. Similarly, the completion of rich behavioural trait datasets for entire taxonomic groups would pave the way towards multi-dimensional community-based analyses of behavioural diversity (BD) metrics, adopting methods from the functional diversity (FD) literature (Petchey & Gaston 2002, Villéger et al. 2008). Setting strategic conservation priorities based on rare behaviours or BD may have important implications for ecosystem function, particularly when focusing on behavioural traits linked to key ecological processes, such as trophic interactions (pollination, seed dispersal, etc.). In addition, there are opportunities for including behaviours in models designed to pinpoint likely future shifts in conservation status by estimating latent extinction risk (Cardillo et al. 2006). The way these models work is to predict threat status for any taxon based on a wide range of attributes and then compare predictions with their observed threat status, thus flagging up any species currently ‘flying under the radar’ (i.e. likely more threated, and thus a higher conservation priority, than indicated by their current conservation status).

### (c) Conclusions

Over recent years, there have been repeated calls for behavioural ecologists to increase their focus on conservation, not least because their study organisms are being driven to extinction by anthropogenic change (Caro & Sherman 2011). Previous authors have suggested that bridging the gulf between these fields might be achieved by applying the experimental or mechanistic approaches predominant in behavioural ecology to conservation research (Linklater 2004), or else returning to more descriptive forms of behavioural ecology potentially relevant to conservation (Caro 2007). However, neither of these approaches are exactly suited to the needs of global conservation assessments which call for simple standardised classifications of basic behavioural traits at ambitious scales, including natural history observations and morphometric measurements. Our analyses show how broad-scale behavioural classifications are now within reach for some major taxa, highlighting the need for continued sampling of basic descriptive information for massive samples of species and pointing the way forward to a deeper integration of the resultant datasets into conservation assessments at the scale of clades, communities and ecosystems.

## Additional Information

### Data Accessibility

Most datasets used in the analyses are openly available in published sources cited in the methods. Where we have used primary data these are provided in the Supplementary Material.

### Authors’ Contributions

J.A.T and A.L.P developed the concepts and compiled data; A.L.P conducted analyses and produced figures; J.A.T wrote the manuscript with substantial input from A.L.P.

## Acknowledgments

We are grateful to Jakob Bro-Jorgenson for inviting us to contribute to this special issue. We thank Stu Butchart for sharing insights into Red List processes, and numerous collaborators and assistants for help with compiling and organizing data, including Monte Neate-Clegg, Ruth Brandt. This research was funded by a Natural Environment Research Grant (NE/I028068/1 to JAT) and a Royal Society University Research Fellowship (to ALP).

