## Supplementary materials for "Integrating behaviour and ecology into global biodiversity conservation strategies"

**­­­­­­**

**Supplementary material**

**Assigning species to dietary niches and foraging behaviours**

Quantitative data on the proportion of resources obtained using different foraging behaviours are generally not available for the majority of species. However, expert descriptions in the literature are usually sufficiently detailed to be translated into coarse categories that capture major differences in foraging behaviours relevant to the global scale of our analysis. Following the method used by Wilman et al. (2014) for quantifying avian diets, we used a standardized protocol to translate qualitative descriptions of foraging behaviour (del Hoyo et al. 2018) into semi-quantitative scores in a systematic way.

For each of the 9 resource types recognised here (aquatic animals, aquatic plants, terrestrial invertebrates, terrestrial vertebrates, terrestrial carrion, nectar, seeds, fruit, other terrestrial plant matter), we scored species for multiple foraging behaviours. In total, across all resource types we recognised 30 different foraging categories, which although not an exhaustive catalogue, reflect the level of detail typically available in published sources and are consistent with previous schemes examining avian guild structure (Fitzpatrick 1985, Croxhall 1987, Remsen & Robinson 1990). For each foraging category, scores were assigned from 0 to 100% in 10% intervals indicating the % of each resource type obtained through that behaviour. In a few very rare cases these scores could be obtained from quantitative information on species foraging strategies. In most cases, however, scores were based on the particular terminology and relative word usage of foraging behaviours described in del Hoyo et al. (2016). For consistency, we aimed to obtain information from this single source but we supplemented this with additional searches of primary literature and online materials where detailed information was lacking. If a single foraging behaviour was described this received a score of 100. Where multiple foraging behaviours were mentioned, we used general terms describing their relative frequency as an initial guide (e.g. ‘mostly’ > 6, ‘sometimes’ = 2, occasionally = 1), adjusting these scores according to the remaining content of the description, family level summary descriptions, as well as additional literature searches. If no indication on the relative use of different behaviours was provided, categories listed earlier in the description were up-weighted relative to those listed at the end. In cases where foraging behaviours were not explicitly mentioned, this information could sometimes be inferred unambiguously from information on the microhabitat and dietary items utilized. By taking the product of the % of the species’ diet consisting of a particular resource type and the % of that resource type obtained through a particular foraging behaviour, we calculated the total % contribution of each of the 30 foraging behaviours to a species’ overall diet.

Some foraging behaviours are unique to a single dietary niche (e.g. bark probing is only utilised by invertivores), while others can be employed for obtaining different resource types (e.g. species foraging on the ground can eat invertebrates, fruits, seeds etc). Because our models predicting threat status contain diet as a factor and in order to limit the number of different foraging behaviours that we estimate an effect for, we distilled the 30 foraging categories into a smaller set of ten categories by lumping analogous behaviours across diets. The final categories we used are: ‘Aerial screen’, ‘Bark glean’, ‘Aerial sally’, ‘Arboreal glean’, ‘Ground forage’, ‘Aquatic ground’, ‘Aquatic plunge’, ‘Aquatic surface’, ‘Aquatic aerial’ and ‘Aquatic dive’.

Based on the scores for each diet category, we assigned species to a single diet category if this contributed to at least 50% of its total diet. If a species scored 50% for two categories, the species was assigned to the category that had the greatest average score across the entire family. This procedure for dealing with tied scores was used because it means that for species to be assigned to a novel category (with respect to other closely related species) there must be a substantial difference in its diet and is thus conservative to errors arising from converting textual descriptions to quantitative scores. If species obtained less than 50% of their resources through any single resources type, the species was classified as an ‘omnivore’, resulting in a total of ten dietary niches. We used an identical protocol to assign each species to a single foraging behaviour. In this case, species employing multiple foraging behaviours in relatively equal proportions were assigned as either ‘aquatic foraging generalists’ or ‘terrestrial foraging generalists’, resulting in a total of twelve foraging behaviour categories.

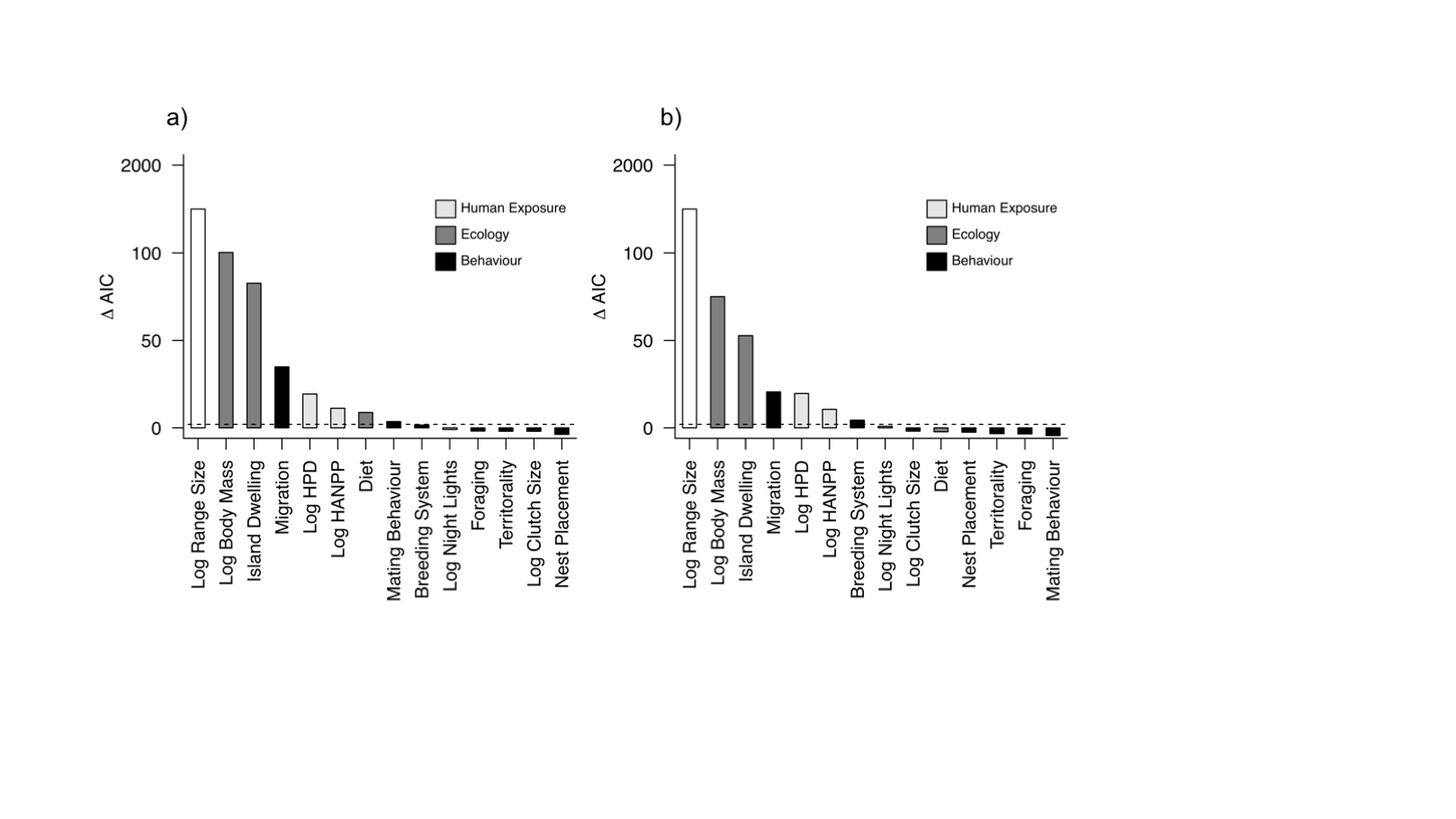

**Figure S1.** The relative contribution of anthropogenic, ecological and behavioural variables to explaining threat status. Results are shown for a) terrestrial (n = 8433) and b) forest dependent species (n = 5646). Variable contributions are quantified as the difference in AIC between the full model and a model excluding each variable. Variables are colored according to variable type. The dashed line indicates a difference of 2 AIC units indicating strong support for variable inclusion.

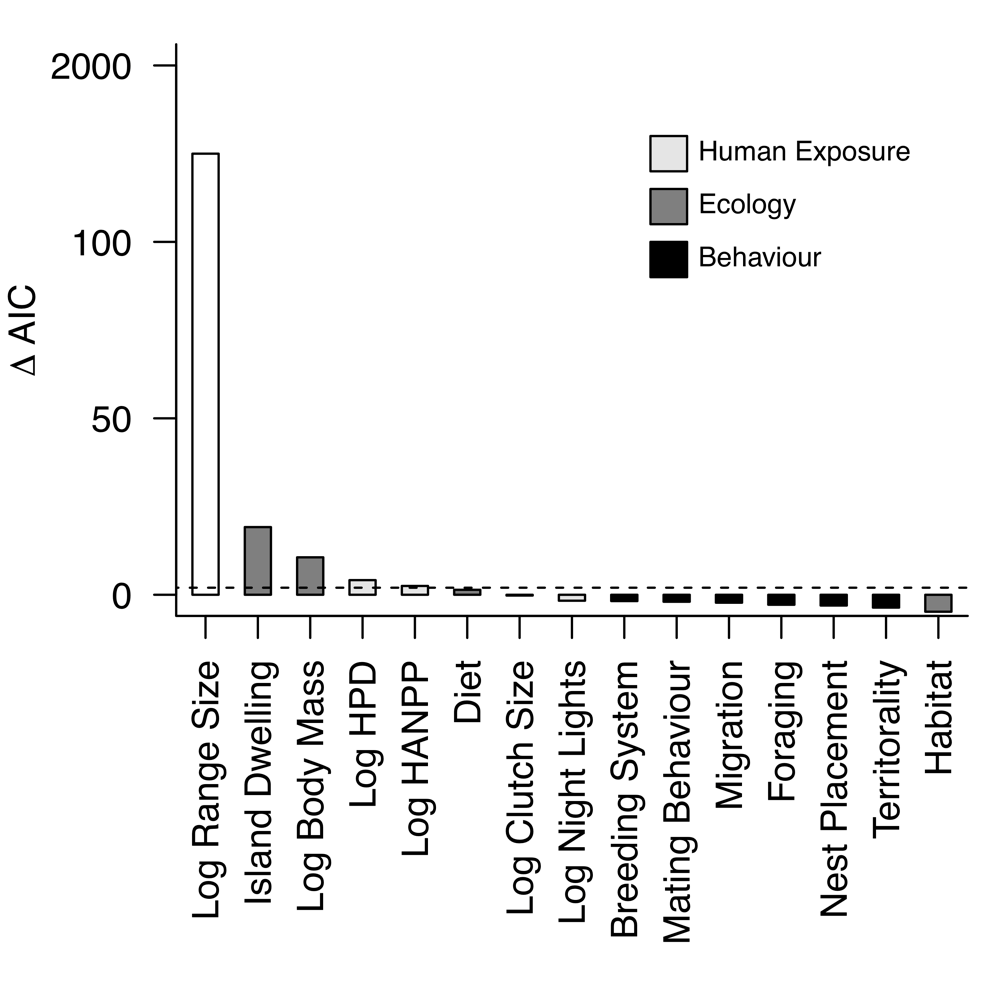

**Figure S2.** The relative contribution of anthropogenic, ecological and behavioural variables to explaining threat level (Vulnerable versus Endangered, Critically Endangered or Extinct). Variable contributions are quantified as the difference in AIC between the full model and a model excluding each variable. Variables are colored according to variable type. The dashed line indicates a difference of 2 AIC units indicating strong support for variable inclusion.

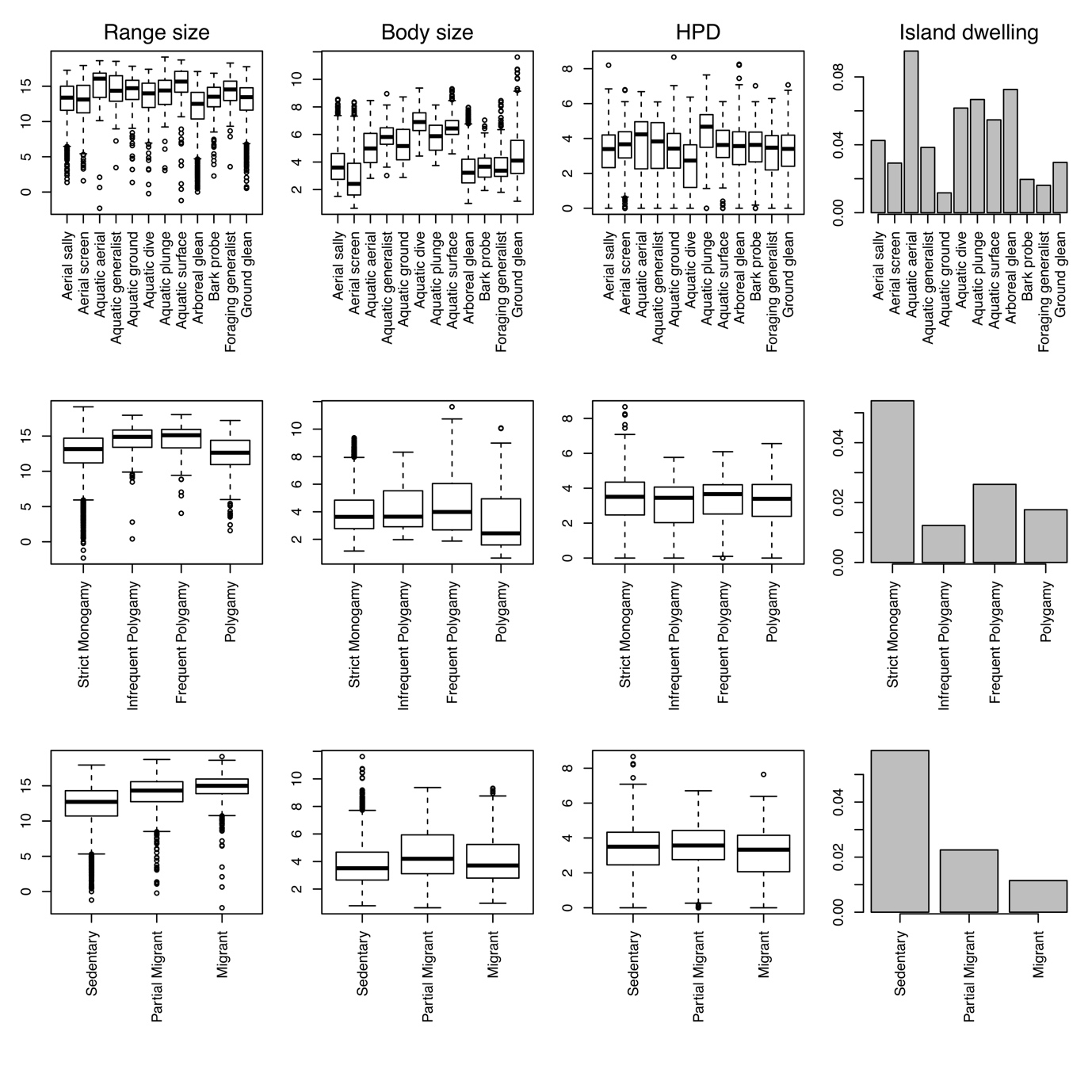

Figure S3. **Associations between behavioural factors (rows) and key drivers of threat status (columns).** Boxplots show the variation in range size (left), body size (middle left), human population density (HPD, middle right) and proportion of island dwelling species (right) across different categories of foraging behaviour (top row), mating behaviour (middle row) and migratory behaviour (bottom row). Range size, body size and HPD are log-transformed.

**Table S1** Predictors of threat across birds (n = 9576). Results are shown for both the full multivariate and individual univariate models. Threat status (0, 1) was modelled using a generalised linear mixed effects model including taxonomic family as a random effect.

|  |  | Multivariate | | | Univariate | | |
| --- | --- | --- | --- | --- | --- | --- | --- |
|  | Variable | Effect | Standard Error | *P* | Effect | Standard Error | *P* |
|  | Intercept | -2.578 | 0.59 | 0 |  |  |  |
|  | LogBodyMass | 0.74 | 0.078 | 0 | 0.455 | 0.046 | 0 |
|  | LogRangeSize | -2.206 | 0.065 | 0 | -1.765 | 0.001 | 0 |
|  | LogNightLights | -0.083 | 0.046 | 0.068 | -0.119 | 0.035 | 0.001 |
|  | LogHumanPopulationDensity | 0.353 | 0.061 | 0 | -0.282 | 0.032 | 0 |
|  | LogHANPP | -0.233 | 0.056 | 0 | -0.425 | 0.033 | 0 |
| Diet | Carrion | 1.232 | 0.695 | 0.077 | 1.037 | 0.457 | 0.023 |
|  | Generalist | -1.462 | 0.489 | 0.003 | -0.541 | 0.185 | 0.003 |
|  | Fruit | -1.34 | 0.504 | 0.008 | -0.356 | 0.196 | 0.069 |
|  | Invertebrates | -1.163 | 0.485 | 0.017 | -0.421 | 0.17 | 0.013 |
|  | Nectar | -1.805 | 0.575 | 0.002 | -0.498 | 0.301 | 0.098 |
|  | AquaticPlants | -1.354 | 0.604 | 0.025 | -0.896 | 0.446 | 0.044 |
|  | TerrestrialPlants | -1.781 | 0.605 | 0.003 | -0.859 | 0.312 | 0.006 |
|  | Seeds | -1.094 | 0.524 | 0.037 | -0.917 | 0.222 | 0 |
|  | Vertebrates | -0.378 | 0.544 | 0.488 | -0.335 | 0.282 | 0.234 |
| Habitat | Sea | -0.631 | 0.453 | 0.164 | 0.318 | 0.308 | 0.302 |
|  | Terrestrial | 0.317 | 0.318 | 0.319 | -0.677 | 0.214 | 0.002 |
|  | Wetland | 0.631 | 0.32 | 0.049 | -0.852 | 0.241 | 0 |
|  | IslandDwellingYes | -1.566 | 0.172 | 0 | 2.102 | 0.114 | 0 |
| Foraging behaviour | AerialScreaning | 0.207 | 0.333 | 0.535 | -0.148 | 0.251 | 0.555 |
|  | AquaticAerial | 1.856 | 0.67 | 0.006 | 1.587 | 0.391 | 0 |
|  | AquaticForagingGeneralist | 0.321 | 0.633 | 0.612 | 1.102 | 0.389 | 0.005 |
|  | AquaticGroundForaging | -0.296 | 0.579 | 0.609 | 0.386 | 0.266 | 0.147 |
|  | AquaticDive | 0.295 | 0.63 | 0.64 | 1.315 | 0.329 | 0 |
|  | AquaticPlunge | 0.002 | 0.776 | 0.998 | 1.265 | 0.469 | 0.007 |
|  | AquaticSurface | 0.445 | 0.567 | 0.433 | 0.805 | 0.325 | 0.013 |
|  | ArborealGleaning | 0.163 | 0.206 | 0.427 | 0.449 | 0.153 | 0.003 |
|  | BarkGleaning | 0.645 | 0.391 | 0.099 | 0.355 | 0.302 | 0.24 |
|  | ForagingGeneralist | -0.439 | 0.574 | 0.444 | -0.99 | 0.523 | 0.058 |
|  | GroundForaging | 0.318 | 0.229 | 0.165 | 0.394 | 0.162 | 0.015 |
| Mating  behaviour | Infrequent Polygyny | 0.572 | 0.366 | 0.119 | -0.518 | 0.29 | 0.074 |
|  | Frequent Polygyny | -0.354 | 0.493 | 0.472 | -0.721 | 0.359 | 0.045 |
|  | Polygamous | 0.643 | 0.238 | 0.007 | 0.167 | 0.181 | 0.355 |
|  | CooperativeBreeding | 0.24 | 0.178 | 0.178 | -0.121 | 0.134 | 0.367 |
| Migration | PartiallyMigratory | 0.383 | 0.161 | 0.017 | -0.478 | 0.123 | 0 |
|  | Migratory | 1.205 | 0.19 | 0 | -0.565 | 0.14 | 0 |
| Territoriality | Strong | -0.259 | 0.178 | 0.145 | -0.054 | 0.127 | 0.668 |
|  | Weak | -0.199 | 0.154 | 0.196 | -0.305 | 0.113 | 0.007 |
| Nest  placement | ExposedElevated | -0.061 | 0.199 | 0.76 | -0.126 | 0.146 | 0.389 |
|  | Cavity | 0.068 | 0.2 | 0.734 | -0.075 | 0.148 | 0.615 |
|  | LogClutchSize | -0.032 | 0.063 | 0.607 | -0.292 | 0.044 | 0 |
